# Polarization of regulatory inputs reveals structural constraints in reconstructed pathway-associated regulatory networks

**DOI:** 10.64898/2026.10.09.758118

**Authors:** Anna Procopio, Rita Granata, Paolo Zaffino, Alessio Merola, Francesco Amato, Carlo Cosentino

## Abstract

Activating and inhibitory interactions jointly shape cellular regulation, yet it remains unclear whether their allocation among individual targets exhibits systematic organization beyond basic structural constraints. Here we introduce antagonistic–cooperative polarization, a target-wise principle in which the sign of each regulatory interaction is determined by the class of its target. This formulation yields a node-separable graph polarization index and decomposes departures from perfect polarization exactly into localized conflicts at targets receiving both activating and inhibitory inputs. We applied the framework to 98 pathway-associated regulatory subnetworks reconstructed from a unified signed regulatory network. Target-wise sign consistency was generally high, but the key evidence for non-random organization came from constrained null models: mixed-input targets were depleted relative to both global sign shuffling and degree-preserving directed rewiring, with depletion increasing systematically with pathway size. This association was robust to rewiring-mobility and pathway-overlap analyses, and indegree-stratified analysis identified the clearest statistically supported depletion among targets receiving exactly two signed inputs. Mixed-input organization was pathway-specific but recurrent across genes; recurrent mixed-input targets remained enriched for Cancer Gene Census genes after accounting for pathway opportunity. Mixed-input occurrences were also preferentially associated with signed feedback motifs, particularly negative feedback, whereas downstream reach was not increased. Together, these results reveal a prevalent polarized backbone punctuated by localized sites of opposing regulation whose abundance departs from expectations under complementary null models controlling for network topology, sign composition, degree structure, and source-specific regulatory output.

## Introduction

Living systems rely on regulatory networks to maintain functional states while responding to changing internal and external conditions. At the cellular level, signaling and transcriptional interactions activate or inhibit their targets, giving rise to directed signed networks that provide a compact representation of regulatory logic. How much can be inferred from this signed topology alone, however, remains an open question. Detailed kinetic models can explain specific systems, but their construction is often limited by uncertain parameters, context-dependent nonlinearities, incomplete measurements, and the necessarily partial nature of reconstructed regulatory networks. A complementary approach is therefore to identify structural constraints encoded by interaction signs while treating their dynamical consequences as conditional on additional model-specific assumptions.

Several lines of work have shown that signed topology can constrain the range of admissible dynamical behaviors. Classical regulatory-circuit theory established necessary sign conditions for specific qualitative behaviors: positive circuits are required for multistationarity, whereas negative circuits are required for sustained oscillations [1–4]. In parallel, monotone dynamical systems theory showed that sign-consistent cooperative or competitive interactions can strongly restrict system behavior [5–8]. Network-motif analyses further demonstrated that local regulatory patterns can be overrepresented relative to randomized networks, indicating that biologically relevant organization may also be concentrated in recurrent local structures [9].

Related graph-theoretical approaches have asked whether biological regulatory networks are close to monotone, balanced, or minimally frustrated organizations [10, 11]. Intracellular regulatory networks have been reported to lie closer to monotone systems than randomized counterparts, and algorithmic approaches have quantified their decomposition into monotone subsystems [12, 13]. Frustration-based approaches have likewise linked signed-network organization to ordered responses, energy-landscape structure, and the ability of coarse-grained regulatory models to reproduce cell-fate behavior [14, 15]. Together, these studies support the view that activating and inhibitory interactions are not distributed arbitrarily. However, they mainly characterize consistency through global node assignments, frustration, monotone decompositions, or relationships between the classes of interacting nodes. Community-polarization formulations, for example, explain edge signs through the joint class membership of their source and target [16, 17]. They do not directly address whether interaction signs are systematically organized according to the individual targets that receive them.

Here we address this question by introducing *antagonistic–cooperative polarization*, a strictly target-determined organization of directed signed networks. In a perfectly polarized network, every node receiving at least one signed input belongs to one of two structural classes: cooperative targets receive exclusively activating inputs, whereas antagonistic targets receive exclusively inhibitory inputs. Thus, the sign of an interaction is determined solely by the class of its target. Here, “cooperative” and “antagonistic” refer to structural input-sign classes rather than dynamical system types. This rule yields a node-separable Graph Polarization Index (GPI), whose optimum assigns each target to the class matching the dominant sign of its incoming regulation. Crucially, departures from perfect polarization are exactly localizable: polarization loss decomposes into target-level sign conflicts occurring exclusively at *mixed-input targets* receiving both activating and inhibitory regulation. The framework therefore converts a global question about signed-network organization into a set of identifiable local deviations.

The same formulation admits an explicitly conditional connection to dynamical modeling. Perfect polarization imposes parity constraints on directed walks and feedback circuits and, if signed edges represent sign-definite off-diagonal regulatory derivatives over the domain of interest, implies a common sign orientation among the represented regulatory inputs entering each state equation. This equation-wise consistency provides a structural bridge to dynamical modeling but does not by itself establish monotonicity, equilibrium existence, stability, or homeostatic behavior, which require additional model-specific assumptions [18, 19]. The algebraic basis and limitations of this connection are detailed in the Supporting Information (SI) Appendix, Section S9.

We applied the framework to 98 pathway-associated regulatory subnetworks reconstructed by projecting curated pathway gene sets onto a unified signed regulatory network. Target-wise sign consistency was generally high, but evidence for non-random organization came from constrained null-model comparisons: mixed-input targets were depleted relative to both global sign shuffling and degree-preserving directed rewiring, with depletion increasing with pathway size. This association remained robust to rewiring-mobility and pathway-overlap analyses, while indegree-stratified analysis localized the clearest statistically supported depletion to targets receiving exactly two signed inputs. Mixed-input organization was pathway-specific but recurrent across genes, and recurrent mixed-input targets remained enriched for COSMIC Cancer Gene Census genes after accounting for pathway opportunity. Mixed-input occurrences were also preferentially associated with signed feedback motifs, particularly negative feedback, whereas greater downstream reach was not observed. Together, these results support an organizing picture in which pathway-associated regulatory subnetworks contain a prevalent polarized backbone punctuated by localized, context-dependent sites of opposing regulation whose abundance departs from constrained null expectations.

## Materials and methods

### Pathway reconstruction and selection

The 98 pathways analyzed in this study were not used as preassembled network objects. Instead, they were reconstructed computationally by combining pathway membership annotations with a global signed and directed gene regulatory network (GRN) downloaded from OmniPath [20]. The reference GRN was assembled from transcription factor–target interactions available through OmniPath. In the executed pipeline, signed regulatory interactions were successfully retrieved from DoRothEA [21] and CollecTRI [22, 23]. DoRothEA interactions were restricted to confidence levels A–C. TRRUST was queried but was not available through the installed OmniPath interface. Interaction endpoints were standardized to HUGO Gene Nomenclature Committee (HGNC) gene symbols, and regulatory sign was inferred from the resource-specific annotation fields, assigning +1 to activating/upregulating interactions and −1 to inhibitory/repressing interactions. After merging all retrieved resources, duplicate edges with identical direction were collapsed. If the same ordered gene pair appeared with discordant signs across resources, that interaction was removed to avoid sign ambiguity. Self-loops were also excluded. This procedure yielded a global signed GRN of 62,340 directed interactions involving 6,983 genes. Pathway membership annotations were then obtained from the OmniPath annotation layer. The code queried pathway-like annotations from the resources NetPath [24], Reactome, KEGG, WikiPathways, SIGNOR [25], ACSN, SignaLink3, PathwayCommons, and MSigDB, retaining annotation records whose label field contained the term “pathway.” Gene symbols were normalized, and duplicate membership entries were removed, resulting in 12,634 pathway-membership records spanning 4,592 annotated genes. Of these, 1,611 genes overlapped with the genes present in the signed GRN. Each candidate pathway was then reconstructed as an induced subnetwork of the global GRN. Specifically, for every annotated pathway, we collected the set of member genes and retained only those GRN interactions whose source and target both belonged to that set. Thus, the analyzed object was not the original pathway annotation itself, but the signed transcriptional regulatory subgraph supported by the OmniPath-derived GRN within that pathway gene set. Candidate pathways yielding no internal GRN interactions were discarded. Consequently, pathway topology was determined entirely by the reconstructed signed GRN rather than by the original interaction structure encoded in the source pathway database. Pathways were retained for analysis only if their reconstructed subnetwork contained at least 3 nodes and at least 2 directed edges. These thresholds were applied to the reconstructed signed subgraph rather than to the raw annotation set. Starting from 127 candidate pathway modules with nontrivial annotation support, this filtering procedure yielded 98 pathways for downstream analysis. All retained interactions had known regulatory signs, so the fraction of sign-resolved edges was effectively 1 in the final analyzed set. The retained pathways originated exclusively from two source databases, SIGNOR signaling-interaction database (*N* = 63) and NetPath (*N* = 35); no pathways from the other queried annotation resources passed all reconstruction and filtering steps. In this framework, each pathway should therefore be understood as a curated gene set from a pathway database, projected onto a unified signed GRN and analyzed only through the subset of directed regulatory interactions that are both internally connected and sign-resolved within that pathway. A stepwise audit of the GRN, pathway-membership table, gene overlap, candidate modules, and final retained subnetworks is provided in SI Appendix, Section S5 and Table S1.

### Graph polarization and optimal polarity assignment

For each pathway, we determined the optimal polarity assignment by maximizing the Graph Polarization Index (GPI), yielding the optimal score *GPI^∗^*. Because the objective depends only on incoming interactions, the optimization is node-separable: the optimal polarity of each target is determined independently by the dominant sign of its incoming edges. The deterministic pseudocode used to compute the optimal polarity assignment, GPI*^∗^*, target-level inconsistency, and mixed-input target fraction is provided in SI Appendix, Algorithm S1. Accordingly,

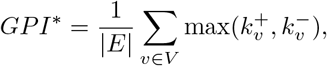

where (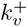) and (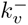) denote the numbers of activating and inhibitory incoming edges of target (v), respectively, and (|.| ) denotes the cardinality. Thus, (*GPI^∗^*) equals the fraction of signed edges agreeing with the dominant incoming sign of their target and quantifies target-wise input-sign consistency. For every graph with at least one signed edge,

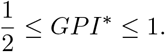

The optimal GPI*^∗^* value is unique, although the corresponding maximizing polarity assignment need not be unique when 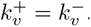. For reproducibility, ties were assigned to the cooperative class *Y* . This deterministic convention does not affect GPI*^∗^*. Nodes with zero signed indegree do not contribute to the objective and may be assigned either polarity without changing the optimal index.

### Inconsistency to polarization and mixed-input targets

Departures from perfect polarization arise exclusively at targets receiving both activating and inhibitory inputs. Let

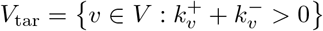

denote the set of nodes receiving at least one signed interaction. A target *v* ∈ *V*_tar_ was classified as mixed-input when

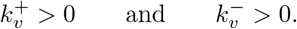

The mixed-input target fraction was defined as

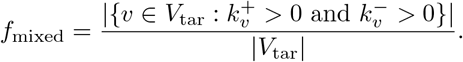

Thus, source-only nodes and nodes with zero signed indegree were not included in the denominator. Node-level inconsistency and its weighted pathway-level average were defined as reported in the Results section.

### Null model construction

To determine whether mixed-input target depletion exceeded that expected from basic topological and sign-related constraints, each reconstructed pathway was compared with ensembles generated under two competitive null models (Table 1). In the global sign-shuffle model, the directed edge set was held fixed, whereas the interaction signs were permuted uniformly among the existing edges. Related fixed-topology sign randomizations have been used to evaluate non-random sign organization in gene regulatory networks. Here, rather than preserving the inhibitory fraction only in expectation, the permutation conditions exactly on the observed pathway-specific numbers of activating and inhibitory interactions. Randomization was performed independently within each pathway. This procedure preserved the pathway-specific global numbers of activating and inhibitory interactions but disrupted their allocation among source–target pairs. In the degree-preserving directed-rewire model, each Markov-chain proposal selected two distinct directed signed edges [26], (*u*_1_*, v*_1_*, s*_1_) and (*u*_2_*, v*_2_*, s*_2_), uniformly from the current edge set and proposed the transformation

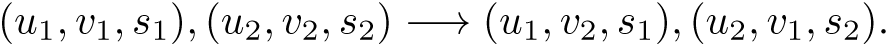

**Table 1.** Structural invariants and inferential roles of the two competitive null models. Both randomizations preserve the numbers of nodes and edges and the pathway-specific global numbers of activating and inhibitory interactions. The signed input signature of a target is 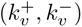, whereas the signed output signature of a source is 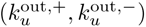. “Exact” denotes node-specific preservation by construction; “not constrained” indicates that the corresponding quantity is not fixed by the randomization.

| Randomization model | Directed topology | Target signed input signature | Source signed output signature | Mixed-input fraction | Inferential role |
| --- | --- | --- | --- | --- | --- |
| Global sign shuffle | Fixed | Not constrained | Not constrained | Not fixed | Topology-conditioned competitive null |
| Degree-preserving directed rewire | Rewired; node-wise in- and out-degree preserved | Not constrained | Exact | Not fixed | Source-constrained competitive null |

A proposal was rejected if the selected edges had the same source or the same target, if the swap generated a self-loop, or if either proposed ordered source–target pair was already occupied. Rejected proposals were retained as Markov-chain self-transitions, rather than resampling until a valid swap was found. This avoids conditioning the proposal probability on the current number of admissible swaps; related detailed-balance and sampling-bias issues in directed degree-preserving randomization are discussed in [27]. Consequently, the degree-preserving procedure preserved exactly the number of edges, the node-wise indegree and outdegree, the pathway-specific global sign counts, and the signed output signature of every source, while allowing the signed input signature of individual targets to vary. It also prevented self-loops, duplicate ordered source–target pairs, and source–target sign conflicts. No realization was discarded or restarted on the basis of its distance from the observed network. All null ensembles were generated independently for each pathway. For each realization, the mixed-input fraction and polarization statistics were recomputed from the randomized network. Pathway-specific null expectations were estimated as ensemble means. Mixed-input target depletion was defined as

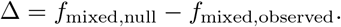

Positive values of Δ therefore indicate fewer mixed-input targets in the reconstructed pathway than expected under the corresponding null model.

### Mobility and proposal-budget diagnostics

Mobility under the implemented directed 2-switch kernel was assessed by exhaustive enumeration of admissible moves from each observed pathway. Pathways with no admissible 2-switch were additionally tested for directed three-cycle reversal and were held fixed if no move was available. For mobile pathways, the proposal budget was calibrated from pilot acceptance rates and evaluated by comparing the null mean mixed-input fraction under baseline and doubled budgets. No pathway exceeded the prespecified budget-sensitivity criterion, and the doubled budget was used for the final ensembles. These diagnostics assess numerical robustness of the focal null statistic and do not establish global state-space connectivity or Markov-chain mixing. Full definitions and results are reported in SI Appendix, Section S7.1-S7.2, Tables S3–S4, Fig. S3, and Supplementary Dataset S3.

### Sensitivity analyses for the degree-preserving null

The primary analysis included all 98 pathways. Sensitivity analyses were repeated in the 76 pathways admitting at least one directed 2-switch and, descriptively, in the 67 pathways whose mixed-input fraction varied across the final null realizations. For each analysis set, we recomputed the Spearman association between depletion and signed-edge count and the one-sided Wilcoxon comparisons for all, small, and large pathways. The size threshold remained fixed at the full-sample median of 32 signed interactions. Complete definitions and results are provided in SI Appendix, Section S7.3.

### Pathway-level mixed-input analysis

Mixed-input targets were identified separately within each pathway, and their frequency and contribution to overall inconsistency were quantified. Pathway-level statistics were computed independently and aggregated without pooling networks. To assess recurrence across biological contexts, nodes were mapped to gene identifiers and tracked across pathways. For each gene, we quantified how often it appears as a mixed-input target relative to its pathway occurrences, together with the associated inconsistency levels.

### Degree-stratified mixed-input analysis

To determine whether mixed-input target depletion was explained solely by singly regulated targets, we repeated the global sign-shuffle analysis after stratifying targets into four disjoint indegree classes:

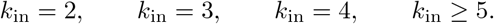

Targets with *k*_in_ = 1 were excluded because they cannot receive both activating and inhibitory inputs by definition. For each pathway and indegree class, the observed mixed-input fraction was compared with the corresponding mean obtained from 1,000 pathway-specific global sign-shuffle realizations. Class-specific depletion was defined as

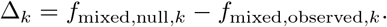

Only pathways containing at least one target in the relevant indegree class contributed to that class-specific analysis. One-sided Wilcoxon signed-rank tests across pathways were used to test whether Δ*_k_* was systematically positive. Because four indegree classes were evaluated, family-wise error was also controlled using the Bonferroni correction.

### Pathway-local downstream reach

For a node *v*, the pathway-local downstream reach score was defined as

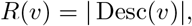

where Desc(*v*) denotes the set of nodes reachable from *v* through directed paths in the reconstructed pathway network. This quantity provides a purely topological measure of pathway-local downstream reach. It does not account for interaction signs, path multiplicity, path length, kinetic parameters, or dynamical state, and should therefore not be interpreted as a direct measure of causal or dynamical influence.

### Gene-level recurrence analysis

To characterize the recurrence of mixed-input organization across biological contexts, target nodes were mapped to gene identifiers and tracked across reconstructed pathways. For each gene, we computed the number of pathways in which it appeared as a signed target, the number and fraction of those target occurrences in which it was classified as mixed-input, and the mean target-level inconsistency and pathway-local regulatory output score across its mixed-input occurrences. This pathway-wise aggregation preserves contextual specificity while enabling the identification of genes that recurrently integrate heterogeneous regulatory inputs. Genes appearing as mixed-input targets in at least two reconstructed pathways were classified as recurrent mixed-input genes. Among recurrent mixed-input genes, the regulator-like subset comprised genes whose mean pathway-local regulatory output score across mixed-input occurrences was greater than or equal to the 75th percentile of the corresponding distribution among recurrent genes.

### Pathway-local regulatory output score

For each reconstructed pathway *P* , let *T_P_* denote the set of signed targets, namely nodes receiving at least one signed incoming interaction within the pathway. For each mixed-input occurrence *v* ∈ *T_P_* , we quantified pathway-local regulatory output as the outgoing degree of the target within the reconstructed pathway, normalized by the maximum outgoing degree observed among signed targets in the same pathway. Formally, the regulatory output score was defined as

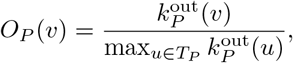

where 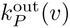 denotes the number of outgoing regulatory interactions from *v* within pathway *P* , and

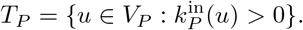

Source-only nodes, defined as nodes with zero signed indegree within the reconstructed pathway, were excluded from the normalization reference set. If the maximum outgoing degree among signed targets in a pathway was zero, the score was set to zero. This quantity was used as a purely topological measure of pathway-local regulatory output and does not incorporate downstream reachability, path length, path multiplicity, interaction sign, or dynamical information.

### Functional enrichment analysis

Functional enrichment was performed separately for the two gene sets derived from the final gene-level recurrence analysis. Recurrent mixed-input genes were defined as genes appearing as mixed-input targets in at least two reconstructed pathways, yielding 117 genes. Among these genes, the regulator-like recurrent subset was defined by a mean pathway-local regulatory output score greater than or equal to the 75th percentile of the corresponding distribution among recurrent mixed-input genes. The resulting cutoff was 0.2387 and identified 30 regulator-like recurrent genes. Enrichment analysis was performed using the gprofiler-official Python client for g:Profiler [28], with *Homo sapiens* as the reference organism. The background universe comprised the 1030 genes appearing as signed targets in the analyzed reconstructed pathways. Gene Ontology biological process, molecular function, and cellular component annotations were tested together with Reactome, KEGG, WikiPathways, CORUM, and transcription-factor target annotations. Statistical significance was assessed using the g:Profiler false-discovery-rate procedure, retaining terms with adjusted *P <* 0.05. Enrichment results were interpreted as descriptive functional annotations and were not used to define mixed-input status, recurrence, regulator-like classification, or pathway polarization.

### Signed feedback motif and degree-matched enrichment analysis

To investigate whether mixed-input targets preferentially localize within structural motifs associated with nonlinear dynamical behaviors, we quantified the signed feedback architecture surrounding each node within every reconstructed pathway. For each pathway-specific signed regulatory graph, we identified directed feedback motifs of length two and three and classified them according to the sign parity of the corresponding cycle. Directed three-node cycles were considered identical only under cyclic rotation of the same ordered cycle. Oppositely oriented directed cycles involving the same three nodes were retained as distinct motifs. Cycles with positive overall sign were interpreted as positive feedback motifs, classically associated with multistability and switching behavior, whereas cycles with negative overall sign were interpreted as negative feedback motifs, classically associated with oscillatory or adaptive dynamics. For every node, we quantified participation in positive and negative directed feedback motifs by counting signed cycles of length two and three within each reconstructed pathway. Positive motifs correspond to cycles with positive overall sign, whereas negative motifs correspond to cycles with negative overall sign. To quantify propagation-related structure, we additionally measured local regulatory output degree and downstream reach, defined respectively as the number of outgoing regulatory interactions and the number of downstream descendants reachable through directed paths. In this analysis, potential for downstream signal propagation was approximated by downstream reach, defined as the number of nodes reachable through directed paths within the reconstructed pathway. To assess structural enrichment while accounting for local connectivity, each mixed node–pathway occurrence was matched to non-mixed nodes from the same reconstructed pathway using indegree and outdegree. Exact degree matching was attempted first; when fewer than three eligible control nodes were available, a common tolerance on indegree and outdegree was progressively increased from 0 to 5. If fewer than three controls remained at the maximum tolerance, all available controls were retained; mixed node–pathway occurrences with no eligible control were excluded. For each mixed node–pathway occurrence, 200 matched control values were sampled with replacement. Enrichment for each metric was expressed as

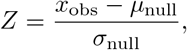

where (*x*_obs_) is the value observed for the mixed-input target and (*µ*_null_) and (*σ*_null_) are the mean and standard deviation of the matched control distribution, respectively. When *σ*_null_ = 0, the enrichment *Z*-score was undefined and was set to missing. One-sided Wilcoxon signed-rank tests were therefore applied only across mixed node–pathway occurrences with finite enrichment *Z*-scores. The downstream-reach metric used in this analysis differs from the pathway-local regulatory output score used in the gene-level recurrence analysis. Whereas downstream reach quantifies the number of descendants reachable through directed paths, the regulatory output score is based solely on normalized local out-degree. For the descriptive gene-level analysis, we retained only gene–pathway occurrences in which the gene was classified as a mixed-input target. For each gene appearing as mixed-input in at least three reconstructed pathways, positive- and negative-feedback motif counts, pathway-local downstream reach, and local regulatory out-degree were averaged exclusively across those mixed-input occurrences. Occurrences in which the same gene was present in a reconstructed pathway but was not classified as mixed-input were excluded from these averages. Point size represents the number of distinct reconstructed pathways contributing a mixed-input occurrence, whereas point color represents mean pathway-local downstream reach across the same occurrences. The gene-level positions were used for descriptive visualization only and were not subjected to a separate enrichment test.

### External cancer-gene annotation analysis

Recurrent mixed-input genes and regulator-like recurrent mixed-input genes were compared with the COSMIC Cancer Gene Census [29]. Gene symbols were standardized to uppercase HGNC-like symbols. The background universe was defined as the set of genes that appeared as signed targets in the reconstructed pathways, thereby restricting the analysis to genes with an opportunity to be detected as mixed-input targets. Enrichment was first tested using one-sided Fisher’s exact tests, followed by Benjamini–Hochberg false-discovery-rate correction. To control for differences in pathway opportunity, we additionally performed an opportunity-matched permutation test in which query genes were randomly sampled from the background within bins defined by the number of pathway opportunities. Empirical *P* values were computed from 10,000 permutations as the fraction of null overlaps greater than or equal to the observed overlap, with a standard plus-one correction. The COSMIC Cancer Gene Census was retrieved using gget [30] from COSMIC version 104, GRCh37, and contained 760 unique cancer genes after gene-symbol standardization. Of these, 246 overlapped the background universe of 1030 signed target genes.

### Computational implementation

The principal pathway-level comparisons used 200 independent realizations per pathway and competitive null model. The final global sign-shuffle and degree-preserving ensembles were generated from master seed 20260702, with separate realization-specific seeds drawn from the corresponding pseudorandom-number generator. The degree-preserving proposal-budget comparison used 20 independent realizations per mobile pathway and proposal budget with seed 20260701. The final degree-preserving ensembles used twice the calibrated baseline proposal budget. The final analysis comprised 19,600 global sign-shuffle realizations and 19,600 degree-preserving realizations. Across all degree-preserving realizations, the number of edges, node-wise indegrees, node-wise outdegrees, pathway-specific global sign counts, and source-specific signed output signatures were preserved exactly. No self-loops, duplicate ordered pairs, or source–target sign conflicts were generated. Degree-stratified global sign-shuffle analyses used 1,000 realizations per pathway with seed 42. Degree-matched feedback-motif analyses used 200 matched control samples per mixed node–pathway occurrence with seed 42. Opportunity-matched COSMIC analyses used 10,000 permutations with seed 123 and a plus-one correction for empirical *P* values.

## Results

### Perfect polarization defines target-wise input-sign consistency

We developed a parameter-independent framework to test whether the signed interactions entering each target in a directed signed network *G* = (*V, E, σ*) exhibit a consistent target-wise organization, as defined in SI Appendix, Section S1 and Fig. S1. Here, *V* is the set of nodes, *E* is the set of directed regulatory interactions, and *σ* : *E* → {+1, −1} assigns an activating (+1) or inhibitory (−1) sign to each interaction.

The framework assigns targets to two structural input-sign classes: a cooperative class *Y* , whose members receive activating inputs, and an antagonistic class *X*, whose members receive inhibitory inputs. The corresponding class-sign function *ε* : *V* → {+1, −1} is defined by *ε*(*v*) = +1 for *v* ∈ *Y* and *ε*(*v*) = −1 for *v* ∈ *X*, where *Y* and *X* denote the cooperative and antagonistic target classes, respectively. Nodes with zero signed indegree do not contribute to polarization and may be assigned to either class. Here, *cooperative* and *antagonistic* denote structural input-sign classes rather than dynamical system types.

In a perfectly polarized network, the sign of each interaction is determined solely by the class of its target, that is,

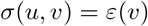

for every directed edge (*u, v*) ∈ *E*. At the graph level, this condition imposes a parity rule on directed walks and feedback circuits, whose signs are determined by the number of antagonistic targets encountered (SI Appendix, Section S3).

Perfect polarization also admits a limited and explicitly conditional connection to dynamical modeling. For a dynamical realization in which signed edges represent sign-definite off-diagonal partial derivatives over the domain of interest, all regulatory derivatives entering the same state equation share a common sign orientation. Thus, derivatives associated with cooperative targets are nonnegative with respect to their represented regulatory inputs, whereas those associated with antagonistic targets are nonpositive. This equation-wise input-sign consistency does not by itself establish monotonicity of the full dynamical system, equilibrium existence, stability, or homeostatic behavior. Its algebraic characterization and limitations are detailed in SI Appendix, Section S9 and Fig. S4.

Across the 98 reconstructed pathway-associated regulatory subnetworks analyzed here (Materials and Methods, Pathway reconstruction and selection; Graph polarization and optimal polarity assignment), optimal Graph Polarization Index (*GPI^∗^*) values were generally high (*GPI^∗^* = 0.91 ± 0.09, mean ± standard deviation; Fig. 1). Thirty-three of 98 subnetworks had *GPI^∗^* ≥ 0.95, including 26 with *GPI^∗^* = 1. Because *GPI^∗^* is a node-separable optimized index with theoretical range [0.5, 1], these raw values are interpreted descriptively as measures of target-wise input-sign consistency. The corresponding computation is formalized in SI Appendix, Section S4 and Algorithm S1. Evidence that the observed target-level sign organization departs from constrained random expectations is evaluated separately below through comparisons of mixed-input target abundance with two null models.

**Fig 1.**
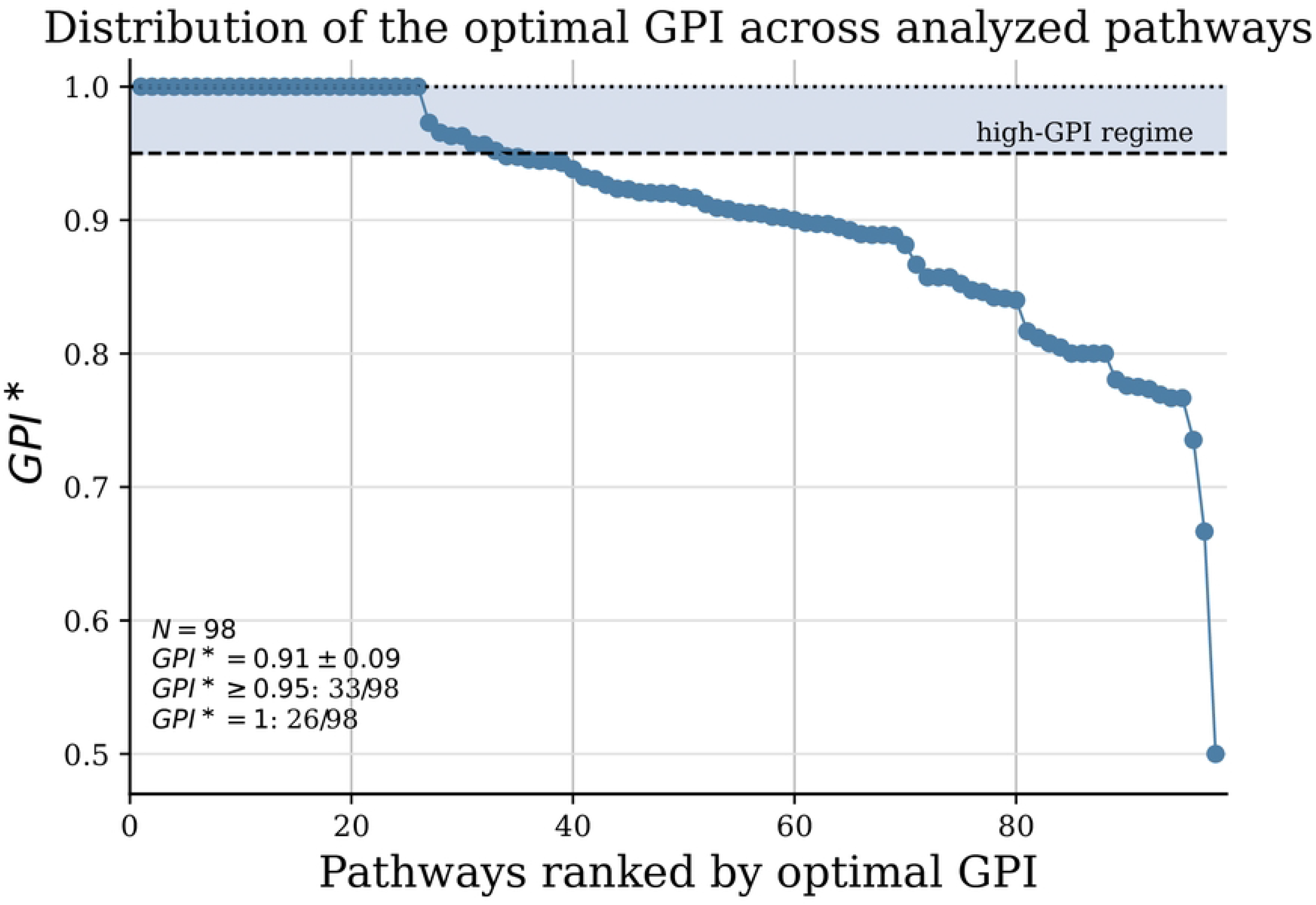
Optimal. *GPI^∗^* **values indicate high target-wise input-sign consistency across the analyzed pathways.** Pathways are ranked by decreasing optimal *GPI^∗^*. Approximately one third of the subnetworks (33/98) had *GPI^∗^* ≥ 0.95, including 26 with *GPI^∗^* = 1.

To illustrate both the limiting case of exact polarization and the localization of departures from it, we examined two representative pathway-associated regulatory subnetworks (Fig. 2). The pentose phosphate pathway attained *GPI^∗^* = 1 and contained no mixed-input targets (Fig. 2A). In this subnetwork, every activating or inhibitory interaction agreed with the sign assigned to its target, yielding complete target-wise input-sign consistency.

**Fig 2.**
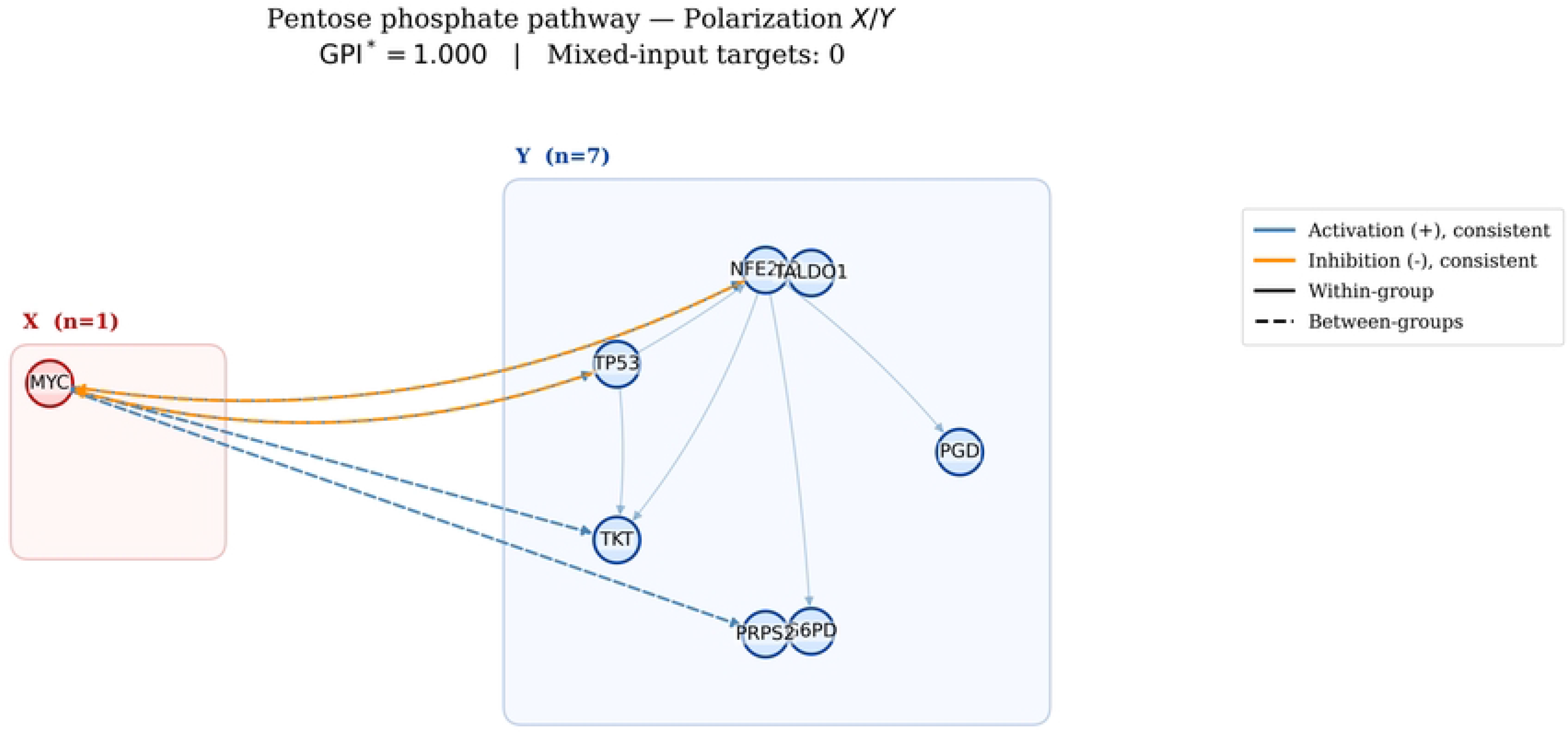
Examples of exact and imperfect target-wise polarization in pathway-associated regulatory subnetworks. (A) The pentose phosphate pathway (*GPI^∗^* = 1.000, and no mixed-input targets). (B) The fibroblast growth factor-1 (FGF1) subnetwork (*GPI^∗^* = 0.889 and 17 mixed-input targets). Blue and orange edges denote activating and inhibitory interactions, respectively; red edges violate the optimal target-wise assignment. Halos identify mixed-input targets, with annotations reporting node-level inconsistency *I_v_*and the number of violating incoming edges. Solid and dashed edges indicate within- and between-class interactions, respectively.

By contrast, the fibroblast growth factor-1 (FGF1) subnetwork attained *GPI^∗^* = 0.889 and contained 17 mixed-input targets (Fig. 2B). The inconsistent interactions were not distributed uniformly throughout the network but converged on a restricted set of targets receiving both activating and inhibitory inputs. This comparison illustrates the node-separable nature of the framework: imperfect global polarization arises from localized sign conflicts superimposed on an otherwise target-wise sign-consistent backbone. Additional representative examples are provided in SI Appendix, Fig. S2.

### Mixed-input targets localize departures from perfect polarization

To identify the structural origin of imperfect polarization, we examined how optimal Graph Polarization Index (*GPI^∗^*) depends on the distribution of incoming interaction signs at target nodes (Materials and Methods, Inconsistency to polarization and mixed-input targets). For each target node *v*, let 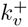 and 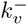 denote the numbers of activating and inhibitory incoming interactions, respectively. Then

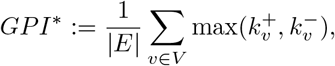

is the fraction of signed edges agreeing with the dominant incoming sign of their target. Departures from perfect consistency therefore arise exclusively at targets receiving mixed-sign inputs.

To quantify this effect, we defined the node-level inconsistency

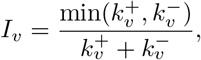

and the corresponding graph-level weighted average

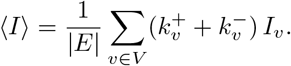

By construction, *GPI^∗^* = 1 − ⟨*I*⟩, showing that the loss of global consistency can be decomposed exactly into local target-level sign conflicts. The formal derivation of the optimized index and its exact decomposition into target-local sign conflicts is provided in SI Appendix, Section S2. This identity is an exact algebraic decomposition of the optimized index and does not, by itself, test whether the observed sign allocation differs from random expectation. Such evidence is obtained from the null-model comparisons presented below. This decomposition indicates that imperfect polarization is not a diffuse property of the network, but is localized to a restricted subset of nodes integrating heterogeneous regulatory inputs. To make this localization explicit, we classified a node as mixed-input when it receives at least one activating and one inhibitory incoming interaction from distinct regulators.

Across pathways, *f*_mixed_ decreased with GPI*^∗^*, consistently with their common dependence on target-level sign conflicts. The two quantities are nevertheless not equivalent: GPI*^∗^* weights conflicts by the number of incident edges, whereas *f*_mixed_ counts the affected targets. A pathway may therefore contain many weakly conflicting targets or a small number of targets accounting for many inconsistent edges. We consequently analyzed *f*_mixed_ explicitly and compared it with constrained null expectations.

To determine whether mixed-input target depletion exceeded that expected from basic topological and sign-related constraints, we compared each reconstructed pathway with two competitive null models (Materials and Methods, Null model construction; Table 1, SI Appendix, Section S7 and Table S2). The global sign-shuffle model preserved the empirical directed topology and the pathway-specific global sign composition while disrupting the allocation of activating and inhibitory signs among target nodes. The degree-preserving directed-rewire model additionally preserved the node-wise indegree and outdegree and the signed output signature of every source, while allowing the signed input composition of individual targets to vary. Together, these competitive models test whether mixed-input target depletion can be explained by global sign composition, degree structure, or regulator-specific output constraints. For each pathway and each competitive null model, we generated 200 independent realizations (Materials and Methods, Computational implementation). The complete pathway-level output table is provided as Supplementary Dataset S1.

Under the implemented degree-preserving kernel, 22 pathways were kernel-immobile and therefore contributed Δ = 0 by construction; the remaining 76 admitted at least one directed 2-switch. The association between mixed-input depletion and pathway size remained significant after restriction to the 76 mobile pathways (Spearman *ρ* = 0.478, *P* = 1.23 × 10*^−^*^5^). Full mobility, proposal-budget, and sensitivity diagnostics are provided in SI Appendix, Section S7, Tables S3–S4, Fig. S3, and Supplementary Datasets S2–S3.

Initial comparisons against randomized controls revealed only a modest average depletion of mixed-input targets across the full pathway collection. However, the effect was highly heterogeneous across pathways. We therefore examined whether the magnitude of mixed-input target depletion depended on pathway size. As a descriptive overview of pathway-level heterogeneity, we also provide a normalized metric atlas summarizing *GPI^∗^*, observed *f*_mixed_, null-model depletion estimates, pathway size, and degree-preserving mobility status across all reconstructed pathways (SI Appendix, Fig. S5).

A clear size-dependent pattern emerged (Fig. 3). Across all 98 pathways, mixed-input target depletion increased with the number of signed interactions under both degree-preserving rewiring (Spearman *ρ* = 0.488, *P* = 3.50 × 10*^−^*^7^) and global sign shuffling (*ρ* = 0.431, *P* = 9.33 × 10*^−^*^6^; Fig. 3A). Similar positive associations were observed when pathway size was quantified by the number of signed targets (degree-preserving rewire: *ρ* = 0.440, *P* = 5.92 × 10*^−^*^6^; global sign shuffle: *ρ* = 0.410, *P* = 2.78 × 10*^−^*^5^).

**Fig 3.**
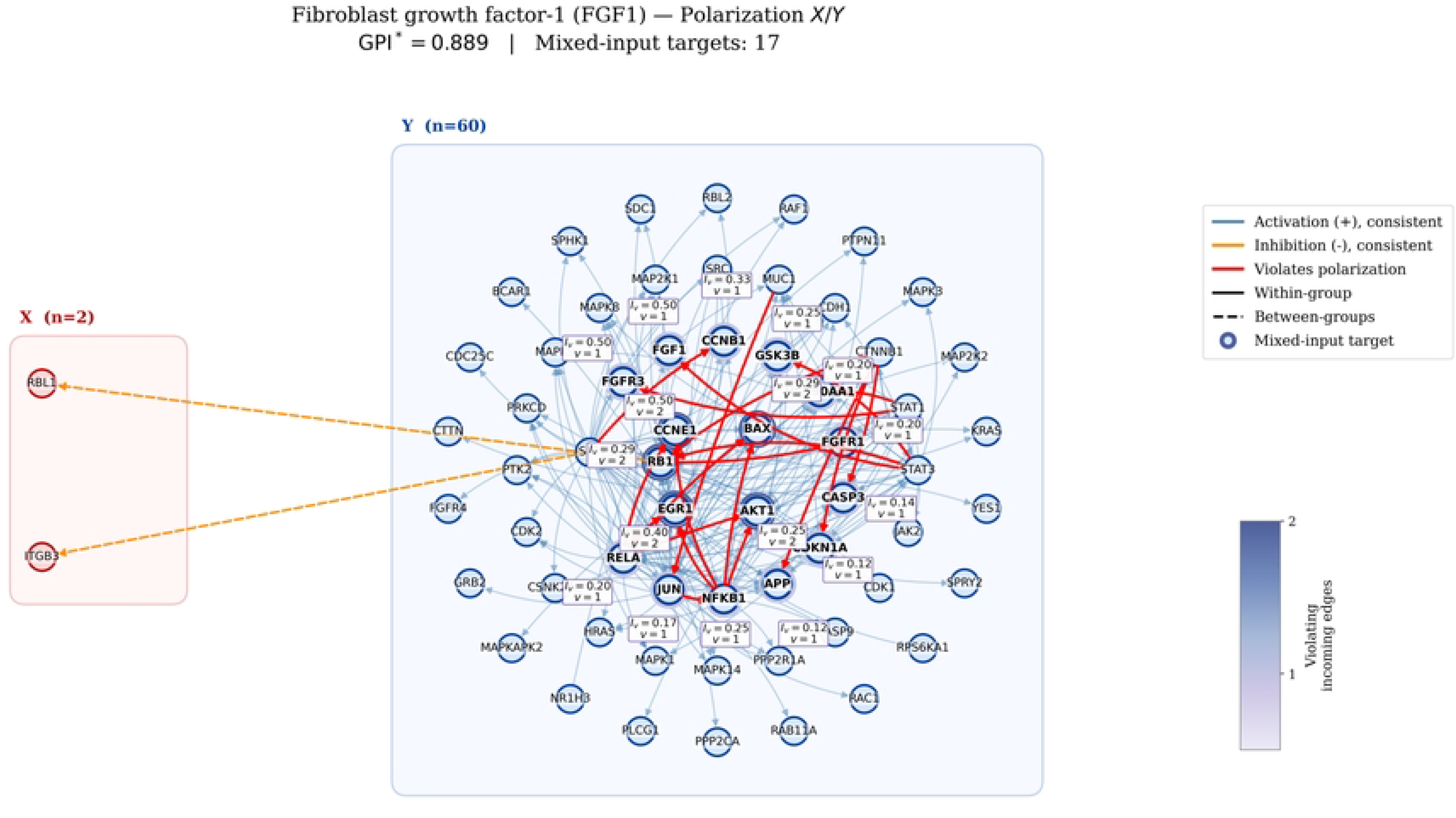
Mixed-input target depletion emerges preferentially in larger pathways. (A) Mixed-input target depletion as a function of pathway size. For each pathway, depletion was defined as Δ = *f*_mixed,null_ − *f*_mixed,observed_, where *f*_mixed,null_ is the mean mixed-input target fraction across randomized networks and *f*_mixed,observed_ is the corresponding fraction in the reconstructed pathway. Positive values therefore indicate fewer mixed-input targets in the observed network than expected under the null model. Solid curves show centered rolling medians computed over windows of 15 pathways; the vertical dotted line indicates the median pathway size of 32 signed interactions. Depletion increased with pathway size under degree-preserving rewiring (Spearman *ρ* = 0.488, *P* = 3.50 × 10*^−^*^7^) and global sign shuffling (*ρ* = 0.431, *P* = 9.33 × 10*^−^*^6^). (B) Depletion in pathways with fewer than 32 signed interactions (small) and at least 32 signed interactions (large). Under degree-preserving rewiring, large pathways had median Δ = 0.0349, with 37 of 49 pathways (75.5%) showing positive depletion (*P* = 6.79 × 10*^−^*^6^). Under global sign shuffling, large pathways had median Δ = 0.0351, with 34 of 49 pathways (69.4%) showing positive depletion (*P* = 3.94 × 10*^−^*^5^). Small pathways showed no significant depletion under either model. All 49 large pathways were mobile under the implemented degree-preserving kernel. (C) Degree-stratified depletion relative to the global sign-shuffle null. Targets with indegree exactly 2 showed significant depletion (*N* = 81 pathways; median Δ = 0.0712, *P* = 5.97 × 10*^−^*^4^; Bonferroni-adjusted *P* = 2.39 × 10*^−^*^3^). The indegree-3, indegree-4, and indegree-≥ 5 classes did not show statistically significant depletion.

By contrast, depletion was not monotonically associated with pathway consistency under either degree-preserving rewiring (*ρ* = −0.140, *P* = 0.169) or global sign shuffling (*ρ* = 0.045, *P* = 0.658). The significant Pearson correlation observed for pathway consistency under global sign shuffling was not supported by the rank-based analysis and was therefore not interpreted as a robust association. These results identify pathway size, rather than global GPI*^∗^*, as the more robust correlate of mixed-input target depletion.

To summarize this size dependence, pathways were divided using the median of 32 signed interactions. Pathways with fewer than 32 signed interactions were classified as small, whereas pathways with at least 32 signed interactions were classified as large (*N* = 49 per group).

Small pathways showed no significant depletion under either competitive null model. Under degree-preserving rewiring, the median depletion was zero (Δ_med_ = 0), with 9 of 49 pathways (18.4%) showing positive depletion (one-sided Wilcoxon signed-rank *P* = 0.148). Under global sign shuffling, the median depletion was also zero, with 13 of 49 pathways (26.5%) showing positive depletion (*P* = 0.309).

In contrast, large pathways showed positive depletion under both null models (Fig. 3B). Under degree-preserving rewiring, the median depletion was Δ_med_ = 0.0349, and 37 of 49 pathways (75.5%) showed positive depletion (one-sided Wilcoxon signed-rank *P* = 6.79 × 10*^−^*^6^). Under global sign shuffling, the median depletion was Δ_med_ = 0.0351, and 34 of 49 pathways (69.4%) showed positive depletion (*P* = 3.94 × 10*^−^*^5^). Pathway-overlap sensitivity preserved this pattern: retaining one medoid per signed-edge Jaccard cluster at *J* ≥ 0.50 (*N* = 92) yielded size–depletion correlations of *ρ* = 0.479 (*P* = 1.35 × 10*^−^*^6^) under degree-preserving rewiring and *ρ* = 0.441 (*P* = 1.07 × 10*^−^*^5^) under global sign shuffling, with consistent results across edge- and gene-based thresholds (SI Appendix, Section S8 and Table S5).

All 49 large pathways were mobile under the implemented degree-preserving kernel.

Conversely, all 22 kernel-immobile pathways belonged to the small-pathway group; excluding them did not alter the null result among the 27 mobile small pathways (Δ_med_ = 0, *P* = 0.148).

Importantly, the depletion was not explained solely by the abundance of singly regulated targets. After assigning targets to disjoint indegree classes, significant depletion remained among targets receiving exactly two signed inputs (*N* = 81 pathways; median Δ = 0.0712; one-sided Wilcoxon signed-rank *P* = 5.97 × 10*^−^*^4^; Fig. 3C; Materials and Methods, Degree-stratified mixed-input analysis). This result remained significant after Bonferroni correction for the four tested indegree classes (*P*_adj_ = 2.39 × 10*^−^*^3^).

Median depletion estimates were also positive for targets with indegree 3 (*N* = 70, Δ_med_ = 0.0198, *P* = 0.102), indegree 4 (*N* = 53, Δ_med_ = 0.0553, *P* = 0.0623), and indegree 5 or greater (*N* = 49, Δ_med_ = 0.0267, *P* = 0.430), but none of these higher-indegree classes reached statistical significance. Complete summary and pathway-level results for all four tested signed-indegree classes are provided in SI Appendix, Supplementary Dataset S4.

Together, the competitive-null and indegree-stratified analyses locate the observed deviation in the target-wise allocation of activating and inhibitory inputs, rather than in global sign composition, degree structure, source-specific signed output, or the abundance of singly regulated targets. A randomization preserving each target’s signed input signature,

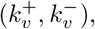

would preserve mixed-input status by construction and would therefore not provide an independent test. The non-random organization was detectable primarily in the larger reconstructed subnetworks.

### Mixed-input targets recur across pathways and frequently coincide with regulator-like nodes

To characterize mixed-input organization without conflating distinct pathway contexts, we analyzed each pathway independently (Materials and Methods, Pathway-level mixed-input analysis; Gene-level recurrence analysis) and identified target nodes receiving at least one activating and one inhibitory input within that pathway. For every mixed-input occurrence, we quantified the associated target-level inconsistency and its pathway-local regulatory output score (Materials and Methods, Pathway-level mixed-input analysis; Pathway-local regulatory output score). We then aggregated these pathway-specific occurrences at the gene level, asking how often the same gene appears as a mixed-input target across its pathway opportunities and whether, when mixed, it also tends to occupy a regulator-like position.

This pathway-wise analysis revealed that mixed-input organization is not uniformly distributed across genes. Most targets were mixed only sporadically or in a restricted subset of their pathway occurrences, whereas a smaller group of genes recurred as mixed-input targets across multiple pathways (Fig. 4A-B). Importantly, recurrence alone did not define a homogeneous structural class. Instead, recurrent mixed-input genes spanned a continuum from local integrators with limited regulatory output score to genes that, when mixed, also displayed high pathway-local regulatory output score, consistent with a regulator-like role. Several genes combined these two properties particularly strongly. Canonical signaling regulators such as *TP53*, *NFKB1*, *MYC*, and *SP1* occupied the high-recurrence, high-regulatory-output region of the gene-level summary, indicating that they are not merely occasional mixed-input targets but recurrent loci in which conflicting upstream regulation coexists with strong pathway-local output connectivity (Fig. 4A). By contrast, other genes showed high mixed-input recurrence but more modest regulatory output scores, suggesting that mixed-input organization can mark either predominantly integrative nodes or more broadly connected regulator-like hubs.

**Fig 4.**
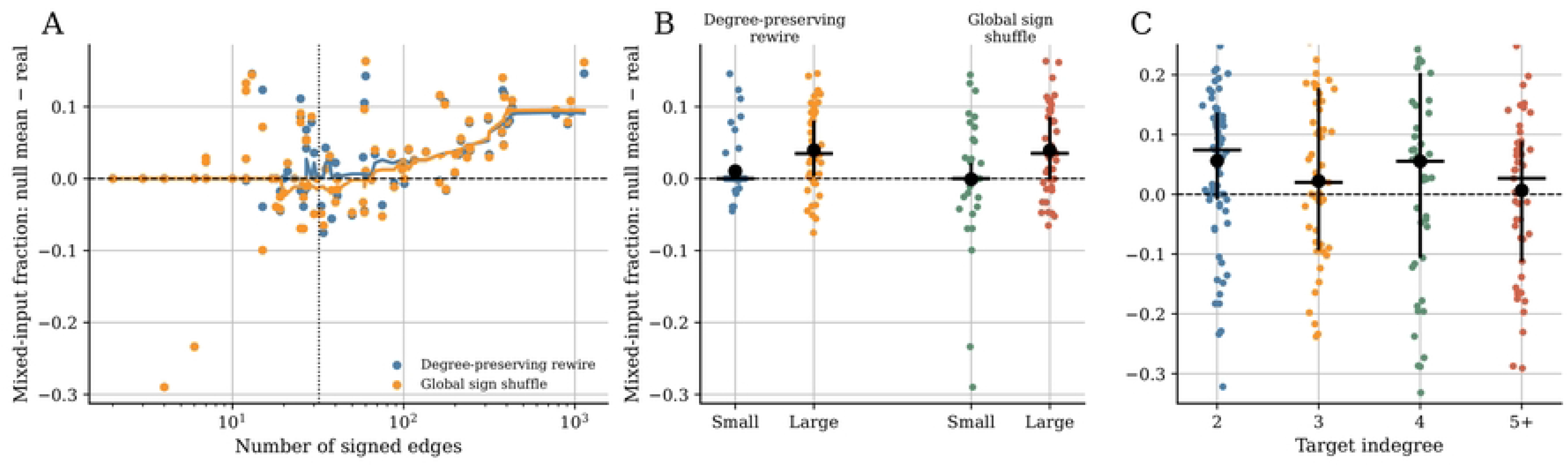
Pathway-wise recurrence of mixed-input targets identifies a subset of regulator-like genes. (A) Gene-level summary of mixed-input recurrence across pathways. Each point represents a gene observed as a mixed-input target in at least one reconstructed pathway and is positioned according to the fraction of pathway opportunities in which it appears as a mixed-input target and its mean pathway-local regulatory output score across mixed occurrences. The regulatory output score is defined as the gene out-degree within the reconstructed pathway, normalized by the maximum out-degree in the same pathway. Point size scales with the number of pathways in which the gene is mixed, and color indicates the mean pathway-local inconsistency across mixed occurrences. Labeled genes highlight recurrent mixed-input targets with prominent regulator-like behavior. (B) Top genes ranked by the number of pathways in which they appear as mixed-input targets. Numbers at the bar ends indicate the fraction of pathway opportunities in which each gene is mixed, distinguishing absolute recurrence from pathway-normalized mixed-input propensity.

Ranking genes by the number of pathways in which they appeared as mixed-input targets identified a recurrent core including *AKT1*, *GSK3B*, *NFKB1*, *RB1*, *JUN*, *MYC*, *STAT3*, *CREB1*, *NFKBIA*, and *TP53* (Fig. 4B). However, absolute recurrence and mixed-input specificity were not identical. For this reason, the fraction of pathway opportunities in which a gene was mixed provides a complementary measure of how consistently mixed-input regulation characterizes that gene across contexts.

Taken together, these results show that mixed-input targets are best understood as a pathway-specific property with gene-level recurrence. Mixed-input organization is not restricted to isolated pathway occurrences, nor is it uniformly associated with strong downstream influence. Rather, a restricted subset of recurrent mixed-input genes repeatedly combines heterogeneous upstream regulation with regulator-like structural features, identifying candidate hubs at which pathway-specific signal integration and local regulatory output connectivity intersect. Complete gene-level recurrence summaries and gene-by-pathway signed-target and mixed-input occurrence data are provided in SI Appendix, Supplementary Datasets S5-S6.

Functional enrichment was analyzed separately for the 117 recurrent mixed-input genes and the 30 regulator-like recurrent mixed-input genes (Materials and Methods, Functional enrichment analysis). The recurrent mixed-input set was strongly enriched for cancer-related regulatory programs, including WikiPathways Cancer pathways (74 genes, false discovery rate (FDR)-adjusted *P* = 4.71 × 10*^−^*^17^), KEGG Pathways in cancer (74 genes, *P* = 6.45 × 10*^−^*^17^), and PI3K–Akt signaling in both WikiPathways (43 genes, *P* = 7.30 × 10*^−^*^11^) and KEGG (44 genes, *P* = 1.18 × 10*^−^*^10^). The regulator-like recurrent subset was characterized by enrichment for transcriptional-regulatory and chromatin-associated functions. Twenty-nine of the 30 genes were annotated with RNA polymerase II cis-regulatory region sequence-specific DNA binding (*P* = 2.05 × 10*^−^*^20^), while all 30 were associated with chromatin (*P* = 2.08 × 10*^−^*^19^) and transcription regulator activity (*P* = 6.17 × 10*^−^*^18^). Because regulator-like classification was based on pathway-local regulatory output within a transcriptional regulatory network, these enrichments are interpreted as functional consistency with the structural classification rather than as independent validation. Complete functional-enrichment results are provided in SI Appendix, Supplementary Dataset S7.

To test whether recurrent mixed-input targets were enriched for independently curated cancer genes, we compared the recurrent mixed-input gene sets against the Catalogue of Somatic Mutations in Cancer (COSMIC) Cancer Gene Census (CGC), using as background the 1030 genes that appeared as signed targets in the reconstructed pathways (Materials and Methods, External cancer-gene annotation analysis). The COSMIC CGC contained 760 genes, of which 246 overlapped the pathway background. Recurrent mixed-input genes showed a strong enrichment for COSMIC CGC genes, with 58 overlaps among 117 recurrent mixed-input genes (Fisher’s exact test, odds ratio (OR) = 3.79, *P* = 9.49 × 10*^−^*^11^, FDR = 1.9 × 10*^−^*^10^). The regulator-like recurrent mixed-input subset showed a similar effect size, with 16 overlaps among 30 genes (OR = 3.83, *P* = 3.87 × 10*^−^*^4^, FDR = 3.87 × 10*^−^*^4^). Because recurrent mixed-input genes may have more pathway opportunities than other genes, we additionally performed an opportunity-matched permutation test. Under this more conservative null model, recurrent mixed-input genes remained significantly enriched for COSMIC CGC membership, with 58 observed overlaps compared with a null expectation of 46.43 overlaps (10,000 permutations; empirical *P* = 0.0046, FDR = 0.0092). By contrast, the regulator-like recurrent subset showed 16 observed overlaps compared with 12.60 expected overlaps and did not remain significant after opportunity matching (empirical *P* = 0.1206). Thus, the COSMIC analysis supports a cancer-gene enrichment of recurrent mixed-input targets overall, while the regulator-like subset should be interpreted as a biologically suggestive, but not independently significant, refinement under opportunity-matched testing.

### Mixed-input targets are enriched in signed feedback motifs

To determine whether mixed-input organization is associated with specific forms of dynamical potential beyond generic network connectivity, we analyzed the signed feedback architecture surrounding mixed-input targets within each pathway. For every node, we quantified participation in positive and negative directed feedback motifs together with pathway-local downstream reach, defined as the number of downstream nodes reachable through directed paths, and local regulatory output (Materials and Methods, Signed feedback motif and degree-matched enrichment analysis; Pathway-local downstream reach).

Under the implemented degree-matching procedure, mixed-input targets showed the strongest enrichment in negative feedback motifs (mean matched *Z* = 1.814, *N* = 149, one-sided Wilcoxon *P* = 7.59 × 10*^−^*^8^). A weaker but significant enrichment was also observed for positive feedback motifs (mean matched *Z* = 0.620, *N* = 181, *P* = 6.74 × 10*^−^*^3^). Local regulatory out-degree retained a smaller positive enrichment (mean matched *Z* = 0.474, *N* = 194, *P* = 4.60 × 10*^−^*^3^), indicating that the matching procedure did not completely eliminate connectivity differences. By contrast, pathway-local downstream reach was not significantly enriched (mean matched *Z* = 0.134, *N* = 101, *P* = 0.701). The feedback-motif associations should therefore be interpreted as partially, rather than fully, independent of local connectivity.

Fig. 5B summarizes genes that appeared as mixed-input targets in at least three reconstructed pathways. For each gene, positive- and negative-feedback motif counts and pathway-local downstream reach were averaged exclusively across pathway occurrences in which that gene was classified as mixed-input. Point size indicates the number of distinct pathways contributing a mixed-input occurrence, whereas color indicates mean downstream reach across the same mixed-input occurrences. Labels are shown for selected recurrent regulators to preserve visual readability. Several labeled genes, including *TP53*, *NFKB1*, *RELA*, *JUN*, *FOS*, *STAT3*, *SP1*, *ESR1*, and *AR*, occupied regions characterized by relatively high mean participation in positive and/or negative feedback motifs when mixed, whereas *MYC* and *HES1* occupied more moderate regions. This panel provides a descriptive gene-level summary and does not constitute a separate enrichment test at the recurrent-gene level. Rather, it identifies pathway contexts in which recurrent mixed-input status co-occurs with feedback-rich local structure. Complete occurrence-level matched-null results and the corresponding node-, gene-, and global-level summaries are provided in SI Appendix, Supplementary Dataset S8.

**Fig 5.**
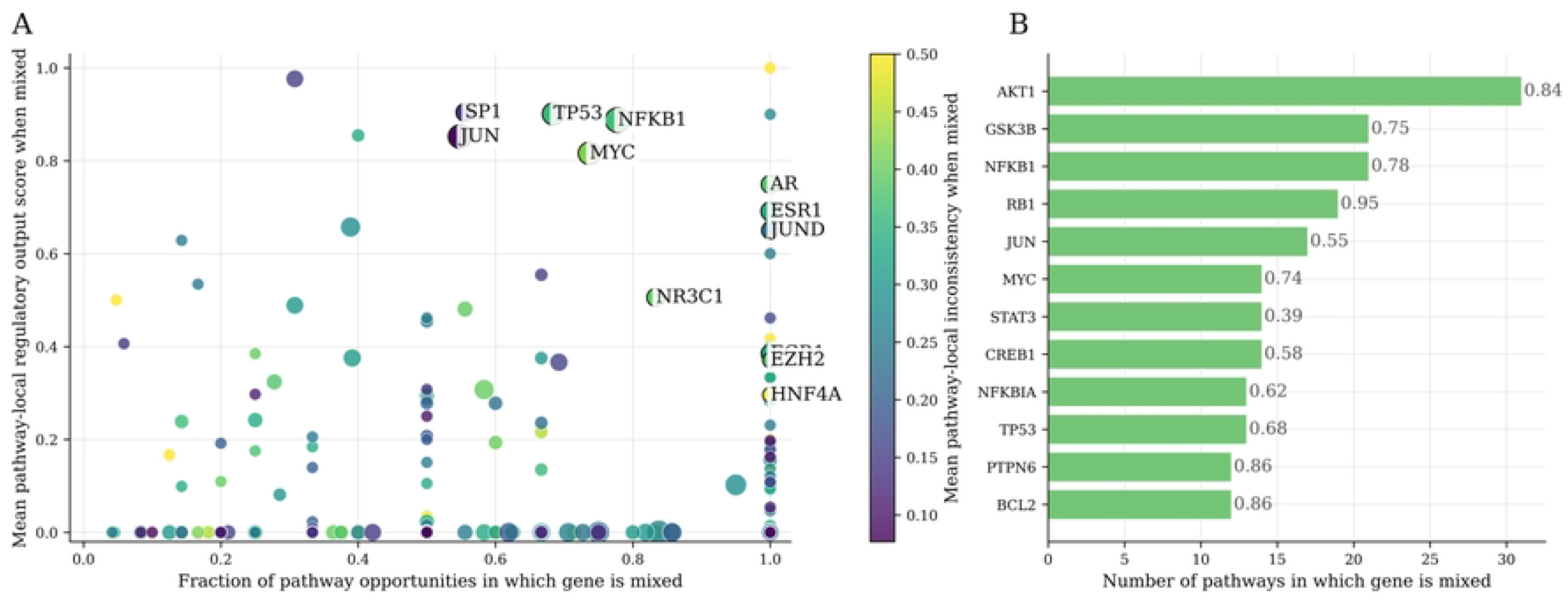
Mixed-input targets are enriched in signed feedback motifs. (A) Degree-matched enrichment analysis comparing mixed-input targets with non-mixed targets from the same reconstructed pathway. Bars indicate mean matched enrichment Z-scores, that denotes the pathway-level standardized difference between the observed mixed-input target statistic and the corresponding degree-matched reference distribution. Mixed-input targets showed the strongest enrichment in negative feedback motifs (mean *Z* = 1.814, *N* = 149, *P* = 7.59 × 10*^−^*^8^), followed by positive feedback motifs (mean *Z* = 0.620, *N* = 181, *P* = 6.74 × 10*^−^*^3^). Local regulatory out-degree retained a smaller positive enrichment (mean *Z* = 0.474, *N* = 194, *P* = 4.60 × 10*^−^*^3^), indicating residual connectivity differences under the implemented matching procedure. Pathway-local downstream reach was not significantly enriched (mean *Z* = 0.134, *N* = 101, *P* = 0.701). P-values correspond to one-sided Wilcoxon signed-rank tests across mixed node–pathway occurrences with finite matched Z-scores. (B) Feedback-motif landscape of genes appearing as mixed-input targets in at least three reconstructed pathways. Each point represents one gene. The horizontal and vertical coordinates indicate the mean numbers of negative and positive feedback motifs, respectively, averaged exclusively across pathway occurrences in which the gene was classified as mixed-input. Point size indicates the number of distinct reconstructed pathways contributing a mixed-input occurrence, and color indicates the mean pathway-local downstream reach across those same mixed-input occurrences.

## Discussion

Antagonistic–cooperative polarization provides a target-centered view of signed regulatory organization by linking a graph-level pattern to the individual nodes that generate its departures. Target-wise consistency was generally high, but raw *GPI^∗^* values were interpreted descriptively because exact polarization can arise more readily in small or sparsely connected subnetworks. Evidence for non-random organization instead came from the depletion of mixed-input targets relative to constrained null models. Depletion became progressively stronger with pathway size and remained evident after accounting for rewiring mobility and pathway overlap (SI Appendix, Supplementary Datasets S1–S3 and S9). After stratification by target indegree, the clearest statistically supported deviation occurred among targets receiving exactly two signed inputs (SI Appendix, Supplementary Dataset S4), indicating that the pattern is already detectable at the simplest level at which activating and inhibitory regulation can converge. Rather than reflecting only global sign composition or generic degree constraints, the observed organization therefore resides in how interaction signs are allocated among individual targets.

A central advantage of the framework is that polarization loss is intrinsically localizable. Most interactions contribute to a broadly target-wise sign-consistent backbone, whereas departures from this organization converge on a restricted set of mixed-input targets. Such localization changes the interpretation of imperfect polarization: sign inconsistency is not treated as a diffuse network property, but as a collection of identifiable target-level conflicts that can be compared across pathways and prioritized individually. Mixed-input status was strongly pathway-dependent, showing that opposing regulation is better understood as a contextual property of a gene within a reconstructed subnetwork than as a fixed gene attribute. Even so, a subset of genes repeatedly appeared as mixed-input targets across distinct pathway settings. After accounting for their opportunities to recur, these genes remained enriched for members of the COSMIC Cancer Gene Census and included prominent regulators such as *TP53*, *NFKB1*, *MYC*, and *SP1* (Fig. 4; SI Appendix, Supplementary Datasets S5–S7). Recurrence thus identifies genes for which heterogeneous upstream regulation repeatedly emerges across biological contexts, while the enrichment for cancer-associated genes provides an independent indication of their potential functional relevance.

The local architecture surrounding mixed-input targets further distinguishes them from ordinary high-degree or broadly influential nodes. After matching for connectivity, mixed-input occurrences remained preferentially associated with signed feedback motifs, with the strongest signal involving negative feedback, whereas downstream reach was not greater than expected (SI Appendix, Supplementary Dataset S8). Their characteristic feature is therefore not enhanced propagation through the reconstructed network, but the convergence of opposing inputs within feedback-rich regulatory neighborhoods. Feedback circuit signs can constrain the repertoire of behaviors available to particular regulatory model classes [1–4]. The observed association places heterogeneous upstream regulation near circuit structures with recognized dynamical relevance and identifies a coherent class of candidate loci for context-specific mechanistic investigation. This distinction is important for experimental design because it favors focused perturbation of individual incoming interactions and pathway-specific input combinations over interpretations based solely on node centrality.

Beyond its biological interpretation, the framework offers a practical computational strategy for analyzing large collections of directed signed networks. The node-separable structure of *GPI^∗^* avoids kinetic parameter estimation and does not require a global combinatorial search, because each target can be evaluated directly from the sign composition of its incoming interactions (SI Appendix, Algorithm S1). The same calculation yields a network-level measure of polarization, identifies the precise targets responsible for polarization loss, and distinguishes the relative balance of opposing inputs from their absolute contribution to the global score. These features make antagonistic–cooperative polarization useful not only for comparing reconstructed pathways, but also for auditing sign annotations, detecting localized inconsistencies, evaluating alternative network reconstructions, and prioritizing targets for mechanistic follow-up. Because the method depends only on directed interaction signs, it can be applied consistently across datasets that differ substantially in size, density, or biological provenance.

Several considerations define the scope of these conclusions. The analysis was intentionally performed on pathway-associated regulatory subnetworks reconstructed through a common projection of curated pathway memberships onto a unified signed regulatory network; the resulting objects should therefore be interpreted as comparable regulatory representations of pathway-associated gene sets rather than as complete biochemical pathway maps. Interaction signs were treated as qualitative regulatory orientations, as required by the graph-theoretic framework, whereas interaction strengths, kinetic parameters, delays, nonlinear response functions, and condition-dependent edge activity were not modeled and would be required to infer specific dynamical behaviors. Twenty-two small pathways admitted no transition under the implemented degree-preserving rewiring kernel and were retained unchanged, contributing zero depletion by construction; however, the size–depletion association remained significant among the 76 mobile pathways, and all large pathways were mobile. Stability under the tested proposal budgets supports the numerical robustness of the focal statistic but is not intended as a formal demonstration of global Markov-chain mixing. Finally, pathway overlap and repeated gene occurrences introduce structured dependence among some observations. The principal pathway-level result remained stable across overlap sensitivities, while opportunity matching accounted for differences in gene recurrence opportunities; nevertheless, the COSMIC and feedback results should be interpreted as complementary structural associations that motivate mechanistic validation rather than as direct evidence of causal or condition-specific regulatory functions.

More broadly, the results support an organizing picture in which pathway-associated regulatory subnetworks are neither uniformly sign-consistent nor randomly conflicted. Instead, they contain a prevalent polarized backbone punctuated by localized, pathway-specific sites of opposing regulation. The reduced abundance of mixed-input targets relative to constrained null expectations indicates that these sites are non-randomly distributed with respect to the structural constraints represented by the implemented null models. Their recurrence across pathways, enrichment for cancer-associated genes, and preferential embedding in feedback motifs further suggest that at least a subset represents biologically meaningful points of regulatory integration. At the same time, the absence of increased downstream reach emphasizes that their relevance is local and organizational rather than a simple consequence of topological dominance.

This combination of global comparison and local interpretability gives the framework its main value. Antagonistic–cooperative polarization converts a broad question about signed-network organization into a tractable set of target-level observations, while preserving information about pathway context, recurrence, feedback embedding, and regulatory output. Applied to context-resolved interaction maps, perturbation-derived networks, or condition-specific reconstructions, the approach could help distinguish persistent sites of opposing regulation from conflicts caused by incomplete annotations or the aggregation of incompatible biological contexts. Integrating these structural predictions with targeted perturbations and mechanistic models should clarify when mixed-input targets act as genuine sites of signal integration, how their roles change across pathways and cellular states, and which of them warrant detailed experimental investigation.

### Data, Materials, and Software Availability

Supplementary Datasets S1–S9 and the Supporting Information Appendix are provided with the article. The code required to reproduce all results reported in the paper is available in a public GitHub repository. Source regulatory interactions and pathway annotations were obtained from OmniPath as described in Materials and Methods. The COSMIC Cancer Gene Census analysis used COSMIC version 104, GRCh37, as described in Materials and Methods.

## Supporting Information

**S1 Appendix. Mathematical definitions, algorithms, and supplementary analyses.** Supporting Information Appendix containing the graph-theoretic formulation of antagonistic–cooperative polarization, derivation of the optimal polarization score and target-level inconsistency measures, algorithmic implementation, dataset-construction audit, null-model and degree-preserving rewiring diagnostics, pathway-overlap sensitivity analyses, and the conditional dynamical interpretation of target-wise polarization.

**S1 Dataset. Pathway-level results.** Complete pathway-level outputs for the 98 reconstructed pathway-associated signed regulatory subnetworks, including observed polarization scores, mixed-input target statistics, null-model summaries, depletion estimates, size-class assignments, and degree-preserving mobility diagnostics.

**S2 Dataset. Null-model definitions.** Machine-readable definitions of the two competitive null models used in the analysis.

**S3 Dataset. Degree-preserving rewire mobility diagnostics.** Pathway-level mobility and rewiring diagnostics for the degree-preserving directed-rewire null model.

**S4 Dataset. Degree-stratified mixed-input depletion results.** Summary statistics and complete pathway-level results for mixed-input target depletion among targets with signed indegree 2, 3, 4, and at least 5.

**S5 Dataset. Gene-level mixed-input recurrence results.** Gene-level recurrence and pathway-local regulatory-output statistics for all signed-target genes, recurrent mixed-input genes, and regulator-like recurrent mixed-input genes.

**S6 Dataset. Gene-by-pathway signed-target and mixed-input occurrences.** Gene-by-pathway signed-target and mixed-input occurrence data in matrix and long formats. Status codes indicate genes that were not signed-target opportunities, signed targets that were not mixed-input, and mixed-input targets.

**S7 Dataset. Functional enrichment and external cancer-gene annotation results.** Complete functional-enrichment results, external cancer-gene annotation analyses, opportunity-matched permutation results, and a compact summary of the enrichment and annotation analyses.

**S8 Dataset. Feedback-motif and degree-matched enrichment results.** Occurrence-level degree-matched null results and corresponding global, gene-level, and node-level summaries for feedback motifs, regulatory out-degree, mixed-input status, and downstream reach.

**S9 Dataset. Pathway-overlap sensitivity analyses.** Complete pathway-overlap sensitivity results, Jaccard-threshold summaries, and signed-edge- and gene-based cluster assignments and representative pathways.

---

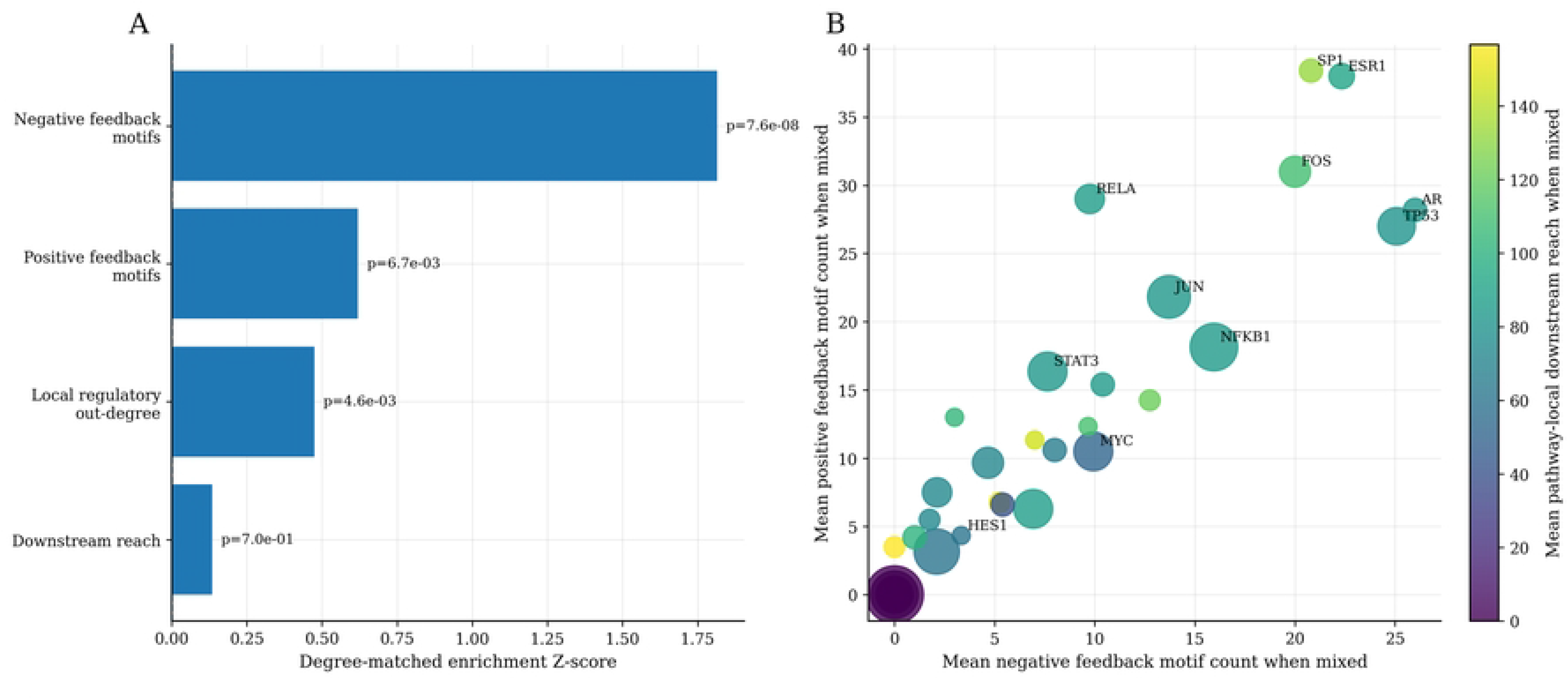

